# Repeated swim exposure and PKN1a knockout enhance group I mGluR-dependent excitability associated with reduced EAAT3 expression in mouse dentate granule cells

**DOI:** 10.64898/2026.08.19.745661

**Authors:** Hiroki Yasuda, Koji Kubouchi, Kenji Hanamura, Taiga Kurihara, Yuka Nakasone, Hideyuki Mukai

## Abstract

Stress-related experiences alter glutamatergic signaling and neuronal excitability, but the mechanisms that couple experience to dentate granule cell function remain incompletely understood. Here, we examined how protein kinase N1a (PKN1a), a protein kinase C-like serine/threonine kinase, and repeated swim exposure regulate mouse hippocampal dentate granule cell excitability, with a focus on the neuronal glutamate transporter excitatory amino acid transporter 3 (EAAT3) and group I metabotropic glutamate receptors (mGluRs). Five days of repeated swim exposure increased spike firing in mature dentate granule cells from wild-type mice. PKN1a knockout produced a similar increase, and repeated swim did not further enhance firing in knockout mice. The enhanced firing observed after repeated swim exposure and in PKN1a knockout mice was reduced by co-application of an mGluR1 antagonist (LY367385) and an mGluR5 antagonist (MPEP). Inhibition of glutamate transporters with DL-TBOA increased granule cell firing in control wild-type mice but did not further increase firing in repeated-swim wild-type or PKN1a knockout mice, suggesting occlusion of transporter-dependent regulation of excitability. Repeated swim exposure and PKN1a knockout also reduced total and surface expression of EAAT3 in the hippocampus, whereas expression of the glial glutamate transporter EAAT2 was not significantly altered. Finally, PKN1a knockout and repeated swim exposure reduced anxiety-related behavior in the elevated plus maze test. Thus, PKN1a-dependent regulation of EAAT3 may restrain group I mGluR-dependent excitability in dentate granule cells, whereas repeated swim exposure and PKN1a knockout shift this system toward a lower-EAAT3, higher-excitability state accompanied by reduced anxiety-related behavior.

## 1. Introduction

Group I metabotropic glutamate receptors (mGluRs), including mGluR1 and mGluR5, are Gq/11-coupled receptors that regulate both synaptic plasticity and neuronal excitability. These receptors induce long-term depression of synaptic transmission (Luscher and Huber, 2010; Malenka and Bear, 2004) and can also enhance neuronal excitation by modulating membrane conductance (Niswender and Conn, 2010). Because mGluR1 and mGluR5 are located perisynaptically in hippocampal neurons (Lujan et al., 1996), their activation is likely to be influenced by local extracellular glutamate levels and the efficiency of glutamate clearance. Therefore, changes in glutamate clearance are likely to influence neuronal excitability through modulation of group I mGluR activation.

The neuronal glutamate transporter excitatory amino acid transporter 3 (EAAT3, also referred to as excitatory amino acid carrier 1, EAAC1) is expressed in hippocampal neurons and can shape glutamate signaling close to neuronal membranes, although its uptake capacity is lower than that of the major glial transporter EAAT2 (Jensen et al., 2015). EAAT3 is colocalized with mGluR5 on neuronal membranes and intracellular organelles in the CA1 region of the hippocampus (Purgert et al., 2014), suggesting that neuronal glutamate transport may influence group I mGluR-dependent signaling. We previously reported that protein kinase N1a (PKN1a), a protein kinase C-like serine/threonine kinase expressed mainly in neurons in the brain, elevates total EAAT3 expression and inhibits group I mGluR-dependent long-term depression (LTD) in the developing hippocampus (Yasuda et al., 2020).

Group I mGluRs have been implicated in several brain disorders, including anxiety and depression (Niswender and Conn, 2010). Systemic administration of group I mGluR antagonists produces antidepressant and anxiolytic effects, which have been hypothesized to result from downregulation of activity in mood– and stress-related brain regions, at least in part through indirect inhibition of NMDA receptors (Niswender and Conn, 2010; Pilc et al., 2008). However, 5 days of chronic restraint stress reduces mGluR5 expression in the CA1 region of the mouse hippocampus and induces anxiety, which is alleviated by local administration of an mGluR5 positive allosteric modulator into the dorsal hippocampal CA1 region (Li et al., 2023). Knockdown of mGluR5 in the CA1 also induces anxiety (Li et al., 2023). Thus, group I mGluRs play a critical role in stress responses (Popoli et al., 2012), and their activity must be tightly regulated in the brain to protect animals from stress-induced dysfunction. However, the mechanisms underlying this regulation remain poorly understood.

Here we focused on the dentate granule cell in the hippocampus, because optogenetic activation of granule cells in the ventral hippocampus reduces anxiety-like behaviors in both open field and elevated plus maze tests (Kheirbek et al., 2013). Furthermore, optogenetic stimulation of hippocampal astrocytes activates dentate granule cells via ATP release and potentiation of synaptic transmission in the dentate gyrus, resulting in reduced anxiety regardless of the dorsoventral hippocampal axis (Cho et al., 2022). Therefore, we examined whether repeated swim exposure, which elevates excitability of pyramidal cells in the cingulate cortex (Sun et al., 2011), influenced excitability of hippocampal dentate granule cells and whether group I mGluRs were involved in changes in granule cell excitability. We also tested involvement of PKN1a in neuronal excitability regulation because PKN1a enhances EAAT3 expression and suppresses group I mGluR-dependent signaling in the hippocampus (Yasuda et al., 2020).

Neonatal maternal separation induces depression but reduces anxiety-related behavior in adolescent rats, in which EAAT3 (EAAC1) expression is reduced in the hippocampus; these behavioral phenotypes are replicated in adolescent EAAT3 KO mice (Kim et al., 2020). In contrast, EAAT3 overexpression in the mouse forebrain prevents depression-like behavior induced by unpredictable chronic mild stress, although anxiety-related behavior is increased even in naïve EAAT3-overexpressing mice (Ardiles et al., 2025; Delgado-Acevedo et al., 2019). These observations also led us to examine whether repeated swim and PKN1a contribute to the regulation of anxiety– and depression-related behaviors as well as dentate granule cell excitability and EAAT3 expression in mice. Here, we show that repeated swim and PKN1a knockout reduce total and surface EAAT3 expression and enhance group I mGluR-dependent firing in mature dentate granule cells. We further show that both manipulations reduce anxiety-related behavior in the elevated plus maze test.

## 2. Material and Methods

### 2.1. Ethics statement

Animal use and all experimental procedures were approved by the Ethical Committee for Animal Experiments of Gunma University Graduate School of Medicine (#9-22, #14-30), the Institutional Animal Care and Use Committee of Kobe University (#17-02-01, #19-5-04), and the Animal Care and Use Committee of Saga University (#30-007-2). All experiments were performed in accordance with the guidelines of these committees.

### 2.2. Repeated swim exposure

PKN1a knockout (KO) mice (Yasuda et al., 2020) and their wild-type littermates at postnatal 8–16 weeks were subjected to repeated swim exposure for 8 min per day for 5 days in a plastic cage (23 × 15 × 16 cm) filled with water to a depth of 10 cm (da Silva et al., 2020; Pericic et al., 2001; Sun et al., 2021). To habituate the mice to water, they were allowed to swim in room-temperature water (22–23°C) on days 1 and 2, during which they adopted a stable head-up, tail-up floating posture. From day 3 onward, they were allowed to swim in 18–20°C water because mice habituated to room-temperature water no longer swam actively. After each swim session, mice were gently dried with a towel. Control wild-type and KO mice were not subjected to repeated swim exposure.

### 2.3. Slice electrophysiology

Hippocampal slices were prepared using a sucrose cutting solution containing (in mM) 215 sucrose, 2.5 KCl, 1.0 CaCl_2_, 4.0 MgSO_4_, 4.0 MgCl_2_, 1.6 NaH_2_PO_4_, 26.2 NaHCO_3_ and 20 glucose, continuously bubbled with 95% O_2_ and 5% CO_2_, as previously described (Zorigt et al., 2025). Slices were first kept for 30–40 min at room temperature in an incubation chamber containing 50% sucrose cutting solution and 50% standard artificial cerebrospinal fluid (ACSF), containing (in mM) 119 NaCl, 2.5 KCl, 4.0 CaCl_2_, 4.0 MgSO_4_, 1.0 NaH_2_PO_4_, 26.2 NaHCO_3_ and 11 glucose. They were then maintained at room temperature in another incubation chamber containing standard ACSF. A slice was transferred to a recording chamber mounted on an upright microscope (BX51WI, Olympus, Tokyo, Japan) equipped with infrared differential interference contrast optics and was perfused with standard ACSF continuously bubbled with 95% O_2_ and 5% CO_2_ at approximately 32°C. Whole-cell recordings were obtained from dentate granule cells using the blind patch-clamp method with recording electrodes filled with an internal solution containing (in mM) 135 potassium gluconate, 10 HEPES, 0.2 EGTA, 10 KCl, 4 Mg-ATP and 0.5 Na_3_GTP (pH adjusted to 7.2 with KOH; osmolarity adjusted to 275–285 mOsm). Membrane potentials were recorded from mature dentate granule cells with membrane resistances greater than 450 MΩ (Goncalves et al., 2016) using a Multiclamp 700A amplifier (Molecular Devices, San Jose, CA, USA). Data acquisition and analysis were performed using custom routines in Igor Pro (WaveMetrics, Portland, OR, USA). Spike firing was monitored in the presence of kynurenic acid (1 mM), an AMPA and NMDA receptor antagonist (Wallace et al., 2009; Weber et al., 2001), or in the presence of CNQX (10–20 μM), D-APV (50 μM) and picrotoxin (100 μM). Voltage responses were recorded in the presence of TTX (1 μM) and Cd^2+^ (200 μM) to assess K^+^ conductance (Wallace et al., 2009). Granule cell spike firing was recorded shortly after break-in to minimize potential time-dependent alterations in firing properties associated with intracellular dialysis during prolonged whole-cell recordings (Ngodup et al., 2024; Pastoll et al., 2012). For each cell, linear regression was applied to the linear portion of the relationship between the number of evoked spikes and the injected-current amplitude (80–280 pA), and the resulting slope was used as an index of neuronal excitability.

D-APV, MPEP, LY367385 and DL-TBOA were purchased from Bio-Techne Tocris Bioscience (Bristol, UK). Other chemicals were obtained from Merck Sigma-Aldrich (Darmstadt, Germany) or FUJIFILM Wako Pure Chemical (Osaka, Japan).

### 2.4. Total protein expression analysis

A mouse was taken from the home cage at the animal facility and immediately anaesthetized with isoflurane. The brain was rapidly removed and placed in ice-cold oxygenated standard ACSF (95% O2/5% CO2). Both hippocampi were dissected, placed in sample tubes and rapidly frozen in liquid nitrogen. Hippocampal lysates were prepared and subjected to immunoblot analysis using an anti-EAAT3 antibody (#12179; Cell Signaling Technology), anti-EAAT2 antibody (#3838, Cell Signaling), and anti-PKN1 antibody(αC6) (Yasuda et al., 2020). Samples were separated by SDS-PAGE and transferred to polyvinylidene difluoride membranes. Membranes were blocked for 1 h at room temperature in PBS (20 mM sodium phosphate, pH 7.5, and 137 mM NaCl) containing 0.05% Triton X-100 (PBS-T) and 5% normal goat serum. Membranes were then incubated for 1 h at room temperature in PBS-T containing the primary antibody. After three washes in PBS-T (5 min each), membranes were incubated for 45 min in PBS-T containing horseradish peroxidase-conjugated secondary antibody at a 1:2000 dilution. Membranes were then washed three times in PBS-T (10 min each), and immunoreactive bands were detected using enhanced chemiluminescence.

### 2.5. Surface EAAT3 expression analysis

Hippocampal slices were washed twice in ice-cold ACSF and incubated with 1 mg/ml NHS-SS-biotin (Pierce) for 30 min at 4°C to biotinylate surface proteins (Nolt et al., 2011). After removal of nonspecifically bound NHS-SS-biotin, tissue was homogenized and sonicated in PBS-based lysis buffer containing (in mM) proteinase and phosphatase inhibitors in 1 EDTA, 1 EGTA, 1% Triton X-100, 0.1% SDS, and 0.1% SDS (pH 7.4), followed by end-over-end rotation for 30 min at 4°C. After centrifugation at 13,000 rpm for 30 min at 4°C, the supernatant was incubated with Neutravidin beads (Thermo Fisher Scientific) to capture biotinylated surface proteins. After three washes with lysis buffer, surface proteins were eluted with protein sample buffer containing DTT and subjected to Western blotting. Samples were separated by 8% SDS-PAGE and transferred to polyvinylidene difluoride membranes. Membranes were blocked for 1 h at room temperature in TBS-T (20 mM Tris/HCl, pH 7.5, 137 mM NaCl and 0.05% Triton X-100) containing 5% Blocking One (Nacalai Tesque). Membranes were then incubated overnight at 4°C in TBS-T containing 5% Blocking One and anti-EAAT3 antibody (Cell Signaling Technology, #12179) at a 1:3000 dilution. After three washes with TBS-T (5 min each), membranes were incubated for 45 min in TBS-T containing 5% Blocking One and horseradish peroxidase-conjugated secondary antibody at a 1:2000 dilution. Membranes were then washed three times in TBS-T (10 min each), and immunoreactive bands were detected using enhanced chemiluminescence.

### 2.6. Behavioral experiments

Male PKN1a KO mice and their wild-type littermates were housed two to three per cage in a room maintained on a 12 h light-dark cycle (lights on at 06.00 h), with ad libitum access to food and water. Before behavioral testing, mice were acclimatized to the experimenters by handling for 3 min per day for 3 consecutive days. Behavioral tests were performed between 09.00 and 18.00 h. To investigate the effects of repeated swim on behavior, mice were subjected to swim for 8 min per session for 5 or 10 consecutive days, as described above, and were gently dried with a towel after each session. Behavioral assessments were conducted 2–5 days after the final swim session. Control mice and mice subjected to repeated swim were tested in the elevated plus maze and tail suspension tests. Only control mice were tested in the forced swim test because mice previously exposed to repeated swim were expected to show altered immobility owing to habituation to water.

#### 2.6.1. Tail suspension test

Mice aged 11–21 weeks were suspended 30 cm above the floor by adhesive tape placed approximately 1 cm from the tip of the tail in a white plastic chamber with a video camera mounted on the side wall (44 cm height × 49 cm length × 32 cm width; O’Hara & Co., Tokyo, Japan). To prevent tail-climbing behavior, the tail was passed through a small plastic cylinder before suspension. Mouse behavior was recorded for 10 min at a sampling rate of 2 frames/s, and immobility was automatically quantified using a predefined threshold in Image FZC software (version 2.20; O’Hara & Co.) (Kojima et al., 2010; Shoji et al., 2022; Steru et al., 1985; Tsuchimine et al., 2020).

#### 2.6.2. Forced swim test

An acrylic cylinder (20 cm height × 10 cm diameter), filled with water at room temperature (23°C) to a depth of 8 cm, was placed in a test chamber with a video camera mounted on the ceiling (49 cm height × 44 cm length × 32 cm width; O’Hara & Co., Tokyo, Japan). Control mice aged 11–21 weeks were placed in the cylinder for 10 min, during which images were captured at a sampling rate of 2 frames/s. Mice were then removed from the cylinder, gently dried and returned to their home cages. The water was replaced between subjects. Immobility during the 10 min session was automatically quantified using Image FZC software (version 2.20; O’Hara & Co.) (Shoji et al., 2022; Tsuchimine et al., 2020).

#### 2.6.3. Elevated plus maze test

The elevated plus maze apparatus consisted of two open arms (25 × 5 cm), two enclosed arms of the same dimensions with transparent walls (15 cm high), and a central square (5 × 5 cm). The apparatus was elevated 50 cm above the floor (O’Hara & Co., Tokyo, Japan). Mice aged 11–14 weeks were placed in the central square, illuminated at 100 lx, facing one of the closed arms. Time spent in the open arms and total distance travelled were recorded for 10 min (Kojima et al., 2010). Data from mice that fell from the maze were excluded from the analyses.

### 2.7. Statistical analysis

Results are reported as means ± SEM. For multiple comparisons, two-way ANOVA or two-way repeated measures ANOVA with Tukey–Kramer test was performed using SigmaPlot version 15 or 16 (Grafiti LLC, Palo Alto, CA, USA) and three-way ANOVA with Šídák’s multiple-comparisons test using EZR (Kanda, 2013). Unless otherwise stated, all other statistical analyses were performed using unpaired, two-tailed Welch’s t tests in Excel.

## 3. Results

### 3.1. Repeated swim exposure and PKN1a knockout enhance group I mGluR-dependent dentate granule cell firing and occlude DL-TBOA-induced excitation

Initially, we investigated whether repeated swim alters neuronal firing in mature dentate granule cells of adult mice and whether PKN1-dependent regulation of group I mGluRs via neuronal glutamate transporters contributes to these changes (Yasuda et al., 2020). Five days of repeated swim did not alter the resting membrane potential and membrane resistance, as assessed by negative current injection at resting membrane potential in both wild-type and PKN1a KO mice (Fig. 1A; control wild-type, *n* = 22 from 6 mice; swim wild-type, *n* = 19 from 3 mice; control KO, *n* = 22 from 4 mice; swim KO, *n* = 30 from 6 mice). However, repeated swim increased spike firing evoked by positive current injection of 200 pA or more in wild-type mice in the presence of kynurenic acid (1 mM), an AMPA and NMDA receptor antagonist (Wallace et al., 2009; Weber et al., 2001), or D-APV (50 μM) and CNQX (10–20 μM), and picrotoxin (100 μM) (Fig. 1B; 200 pA, *p* = 0.014; 240 pA, *p* = 0.001; 280 pA, *p* < 0.001; 320 pA, *p* < 0.001; 360 pA, *p* < 0.001; 400 pA, *p* < 0.001; effects of swim, *F*_(1, 34)_ = 10.281, *p* = 0.003; Two-way repeated measures ANOVA with Tukey–Kramer test). However, repeated swim did not affect spike firing in KO mice (Fig. 1B; effects of swim, *F*_(1, 50)_ = 0.590, *p* = 0.446; Two-way repeated measures ANOVA). We also analyzed the neuronal excitability by calculating slopes of current amplitude-spike firing relationship of each cell and found that repeated swim elevated spike firing (Fig. 1C; control, 0.0420 ± 0.0038 spikes/pA; swim, 0.0730 ± 0.0061 spikes/pA, *p* = 0.002; interaction between swim and genotype, *F*_(1, 89)_ = 8.142, *p* = 0.005; two-way ANOVA with Tukey–Kramer test). PKN1a knockout likewise increased firing in the absence of repeated swim compared with control wild-type mice, and repeated swim did not further increase excitability in KO mice (Fig. 1 C; control, 0.0730 ± 0.0062 spikes/pA, significantly higher than control wild-type at *p* = 0.001, two-way ANOVA with Tukey–Kramer test; swim, 0.0673 ± 0.0073 spikes/pA), suggesting that repeated swim and PKN1a knockout engage overlapping mechanisms to enhance neuronal excitability.

**Figure 1.**
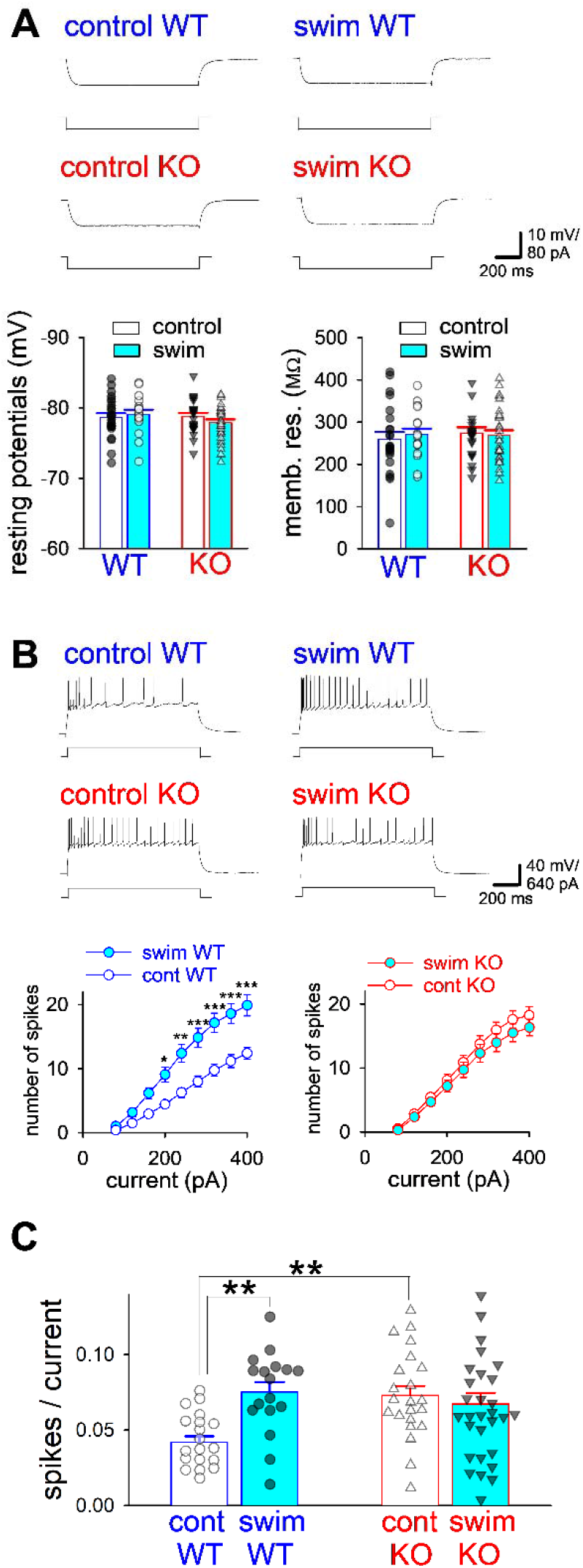
Repeated swim and PKN1a knockout increase spike firing in mature hippocampal dentate granule cells. (A) Membrane hyperpolarization induced by negative current injection (−40 pA, 1 s) (upper), resting membrane potentials (lower left), and membrane resistance at resting membrane potential (lower right) in mature dentate granule cells from control and repeated-swim wild-type (control, *n* = 22 from 6 mice; swim, *n* = 19 from 3 mice) and PKN1a KO mice (control, *n* = 22 from 4 mice; swim, *n* = 30 from 6 mice). Resting membrane potentials and membrane resistances did not differ between genotypes or treatments. (B) Representative spike firing traces (upper) and summary of the injected current–spike number relationship (lower) in mature dentate granule cells from control and repeated-swim wild-type and PKN1a KO mice. (C) Summary of averaged slopes of the injected current–spike number relationship in control and repeated-swim wild-type and PKN1a KO mice. Slopes were calculated using data points within the linear range of the relationship between injected current amplitude and spike number; data points showing saturation at higher current amplitudes, if any, were excluded from the analysis. \**p* < 0.05, **, *p* < 0.01, \*\*\**p* < 0.001. All data are shown as means ± s.e.m.

We first examined whether repeated swim alters spike width during current injection, because spike width reflects Ca^2+^-dependent K^+^ channel activity (Brenner et al., 2005; Sun et al., 2021). The half-widths of the first seven spikes in control wild-type neurons, in which a 1 s, 400 pA current injection evoked approximately 15–16 spikes (Fig. S1A and B; 15.9 ± 1.3 spikes, *n* = 8 from 2 mice), were similar to those in repeated-swim wild-type neurons, in which a smaller current injection evoked a comparable number of spikes (Fig. S1 A and B; 15.9 ± 0.7 spikes, mean current amplitude 302.2 ± 19.0 pA, *n* = 9 from 2 mice). By contrast, the half-widths of the last three spikes were significantly smaller in repeated-swim wild-type mice than in control wild-type mice (Fig. S1A, B, and D; significant main effects of experimental condition (*F*_(4, 32)_ = 8.202, *p* < 0.001) and spike position (*F*_(2, 64)_ = 3.381, *p* = 0.040), as well as a significant condition × spike position interaction (*F*_(8, 64)_ = 2.561, *p* = 0.017), Two-way repeated-measures ANOVA; third from last, control, 1.92 ± 0.10 ms; swim, 1.54 ± 0.04 ms; *p* = 0.005; second from last, control, 1.91 ± 0.09 ms; swim, 1.56 ± 0.05 ms; *p* = 0.011; last, control, 1.94 ± 0.10 ms; swim, 1.59 ± 0.05 ms; *p* = 0.009, post hoc Tukey–Kramer test). However, when spikes were evoked by the same 400 pA current injection, the half-widths of the last three spikes were similar between repeated-swim and control wild-type mice (Fig. S1A, B, and D). If the enhanced firing in repeated-swim wild-type mice were due to reduced Ca^2+^-dependent K^+^ channel activity, spike width would be expected to increase. This was not observed, suggesting that the enhanced firing in repeated-swim wild-type mice is unlikely to be explained by reduced Ca^2+^-dependent K^+^ channel activity.

In KO mice, comparable numbers of spikes were evoked by current injections of approximately 233 pA and 250 pA in control and repeated-swim groups, respectively, and repeated swim did not broaden spike half-width (Fig. S1C and D; control, 15.8 ± 0.3 spikes, 233.3 ± 21.7 pA, *n* = 6 from 3 mice; repeated swim, 15.9 ± 0.4 spikes, 250.0 ± 12.5 pA, n = 8 from 3 mice), suggesting that repeated swim also does not alter Ca^2+^-dependent K^+^ channel activity in KO mice. We next examined the possible involvement of voltage-dependent K^+^ channels by recording voltage responses to current injection in the presence of Na+ and Ca^2+^ channel inhibitors (Dong et al., 2006; Wallace et al., 2009). The amplitudes of the voltage responses did not differ between control and repeated-swim wild-type or PKN1a KO mice (Fig. S1E), suggesting that membrane resistance mediated by K^+^ conductance is not altered (Dong et al., 2006; Wallace et al., 2009). The amplitudes of the voltage responses did not differ between control and repeated-swim wild-type (Fig. S1E; control, *n =* 11 from 2 mice; swim, *n =* 9 from 2 mice) or PKN1a KO mice (Fig. S1E; control, n = 11 from 2 mice; swim, n *=* 16 from 2 mice), suggesting that membrane resistance mediated by K^+^ conductance is not altered (Dong et al., 2006; Wallace et al., 2009).

We next examined the involvement of group I mGluRs, because dentate granule cells express mGluR1 and mGluR5 (Lujan et al., 1996), these receptors enhance neuronal excitation (Niswender and Conn, 2010), and we previously reported that LTD dependent on both mGluR1 and mGluR5 is induced in immature PKN1a KO mice (Yasuda et al., 2020). Although group I mGluR antagonists MPEP (10 μM) and LY367385 (LY; 100 μM) did not consistently alter resting membrane potential (Fig. 2A), they significantly reduced membrane resistance (Fig. 2A; main effect of drug, *F*_(1, 115)_ = 29.551, *p* = 3.10 × 10; three-way ANOVA). Šídák-adjusted comparisons showed no significant difference in the control WT group (Fig. 2A; no drug, 239.7 ± 19.3 MΩ, n = 6 cells from 2 mice; MPEP + LY, 176.5 ± 14.9 MΩ, n = 10 cells from 2 mice; *p* = 0.194). In contrast, MPEP + LY significantly reduced membrane resistance in the swim WT group (Fig. 2A; no drug, 268.2 ± 11.0 MΩ, n = 29 cells from 6 mice; MPEP + LY, 202.4 ± 16.8 MΩ, n = 15 cells from 6 mice; *p* = 0.0049), the control KO group (Fig. 2A; no drug, 278.0 ± 21.8 MΩ, n = 10 cells from 3 mice; MPEP + LY, 196.6 ± 13.4 MΩ, n = 15 cells from 3 mice; *p* = 0.0072), and the swim KO group (Fig. 2A; no drug, 266.1 ± 17.1 MΩ, n = 18 cells from 5 mice; MPEP + LY, 204.9 ± 15.4 MΩ, n = 20 cells from 5 mice; *p* = 0.0126; Šídák’s multiple-comparisons test).

**Figure 2.**
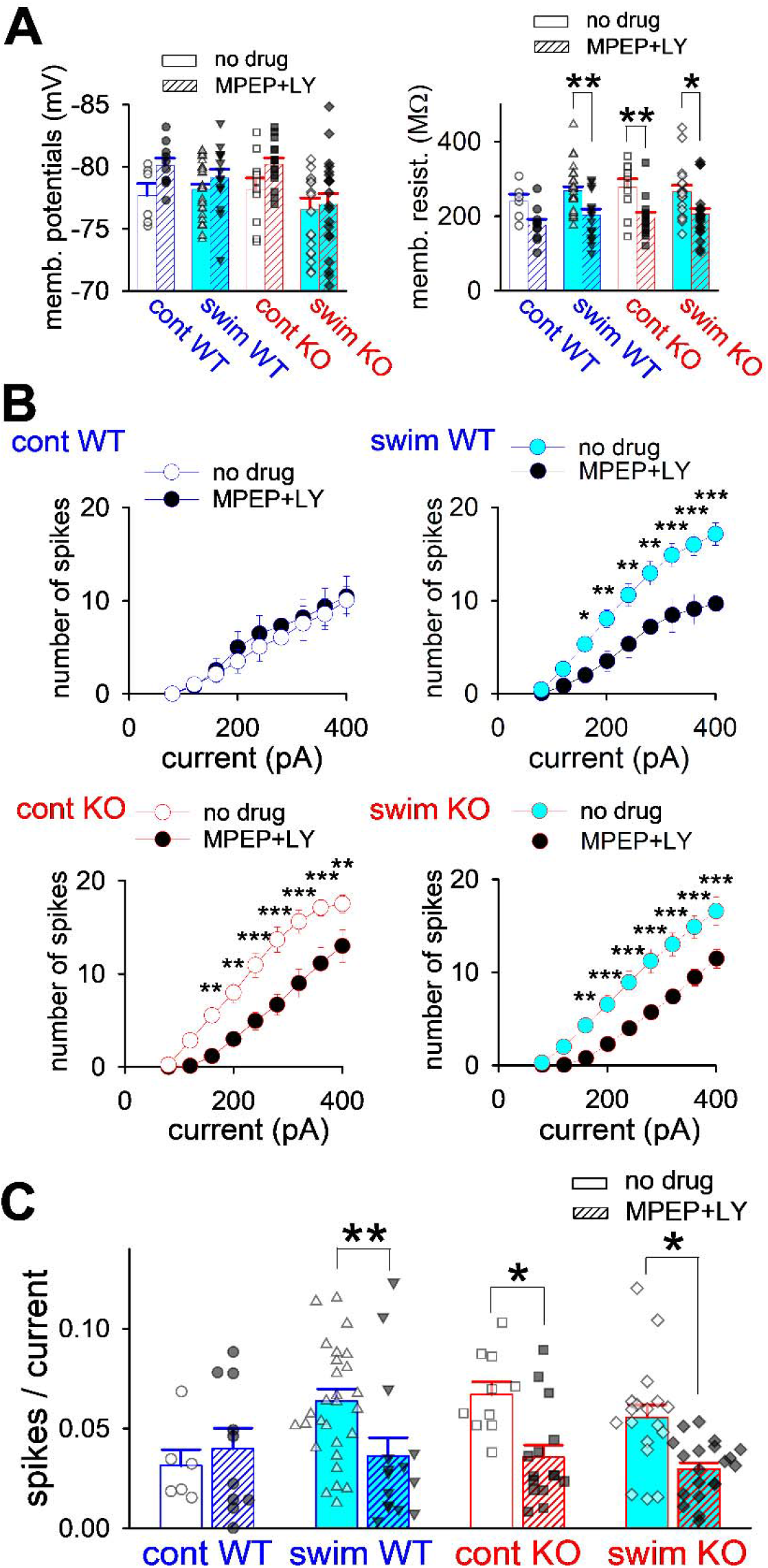
Increased spike firing in repeated-swim and PKN1a KO mice is group I mGluR-dependent. (A) Summary of the effects of group I mGluR antagonists (MPEP 10 μM + LY367385 (LY) 100 μM) on resting membrane potentials and membrane resistance (control wild-type, no drug, *n* = 6 from 2 mice, MPEP + LY, *n* = 10 from 2 mice; swim wild-type, no drug, *n* = 29 from 6 mice, MPEP + LY, *n* = 15 from 6 mice; control KO, no drug, *n* = 10 from 3 mice, MPEP + LY, *n* = 15 from 3 mice; swim KO, no drug, *n* = 18 from 5 mice, MPEP + LY, *n* = 20 from 5 mice). Inhibition of group I mGluRs reduced membrane resistance irrespective of genotype or treatment, suggesting that group I mGluRs regulate excitability even at resting membrane potentials. (B and C) Summary of the effects of group I mGluR antagonists on current-spike relationship (B) and averaged slopes (C) in control and repeated-swim wild-type and PKN1a KO mice. \**p* < 0.05, \*\**p* < 0.01, \*\*\**p* < 0.001. All data are shown as means ± s.e.m.

Group I mGluR antagonists did not affect firing in control wild-type mice (Fig. 2B). However, they reduced firing in repeated-swim wild-type (Fig 2B; drug, *F*_(1, 42)_ = 10.870, *p* = 0.002; drug × injected-current amplitude interaction, *F*_(8, 336)_ = 6.462, *p* < 0.001; 160 pA, *p* = 0.049; 200 pA, *p* = 0.005; 240 pA, *p* = 0.002; 280 pA, *p* = 0.001; 320 pA, *p* < 0.001; 360 pA, *p* < 0.001; 400 pA, *p* < 0.001; two-way repeated-measures ANOVA), control KO (Fig. 2B; drug, *F*_(1, 23)_ = 13.613, *p* = 0.001; drug × injected-current amplitude interaction, *F*_(8, 184)_ = 4.583, *p* < 0.001; 160 pA, *p* = 0.008; 200 pA, *p* = 0.003; 240 pA, *p* < 0.001; 280 pA, *p* < 0.001; 320 pA, *p* < 0.001; 360 pA, *p* < 0.001; 400 pA, *p* = 0.006; two-way repeated-measures ANOVA) and repeated-swim KO mice (Fig. 2B; drug: *F*_(1, 36)_ = 18.922, *p* < 0.001; drug × injected-current amplitude interaction: *F*_(8, 288)_ = 6.110, *p* < 0.001; 160 pA, *p* = 0.004; 200 pA, *p* < 0.001; 240 pA, *p* < 0.001; 280 pA, *p* < 0.001; 320 pA, *p* < 0.001; 360 pA, *p* < 0.001; 400 pA, *p* < 0.001; two-way repeated-measures ANOVA). We also calculated the slope of the relationship between the number of evoked spikes and the injected-current amplitude and found that they significantly reduced the slope (Fig. 2C; drug, *F*_(1, 115)_ = 11.907, *p* = 7.83 × 10; three-way ANOVA). Although these antagonists did not change the slope in control wild-type mice (Fig. 2C; no drug, 0.0313 ± 0.0081 spikes/pA; MPEP + LY, 0.0399 ± 0.0102 spikes/pA), they reduced the slope in repeated-swim wild-type mice (Fig. 2C; no drug, 0.0636 ± 0.0061 spikes/pA, MPEP + LY, 0.0361 ± 0.0093 spikes/pA; *p* = 0.0089, Šídák’s multiple-comparisons test), and control and repeated-swim KO mice (Fig. 2C; control no drug, 0.0595 ± 0.0029 spikes/pA, control MPEP + LY, 0.0435 ± 0.0061 spikes/pA, *p* = 0.0240, Šídák’s multiple-comparisons test; swim no drug, 0.0524 ± 0.0045 spikes/pA; swim MPEP + LY, 0.0377 ± 0.0033 spikes/pA, *p* = 0.0192). These results suggest that the elevated firing induced by repeated swim and PKN1a knockout is mediated by group I mGluRs.

Previously, we reported that PKN1a upregulates expression of the neuronal glutamate transporter EAAT3, and that the glutamate transporter inhibitor DL-TBOA (TBOA) mimics the heterosynaptic LTD observed in PKN1a KO mice (Yasuda et al., 2020). To determine whether this effect involves altered intrinsic membrane properties, we examined the effects of TBOA on resting membrane potential and membrane resistance. In contrast to group I mGluR antagonists, which significantly reduced membrane resistance (Fig. 2A), TBOA (100 μM) did not affect either resting membrane potential nor membrane resistance under any condition tested (Fig. 3A; control wild-type, no drug, *n* = 10 from 3 mice, TBOA, *n* = 10 from 3 mice; swim wild-type, no drug, *n* = 11, from 3 mice, TBOA, *n* = 12 from 3 mice; control KO, no drug, *n* = 9 from 3 mice, TBOA, *n* = 11 from 3 mice; swim KO, no drug, *n* = 18 from 3 mice, TBOA, *n* = 19 from 3 mice). These findings suggest that low levels of ambient glutamate tonically activate group I mGluRs in mature dentate granule cells, even though glutamate transporters are largely ineffective at resting membrane potential.

**Figure 3.**
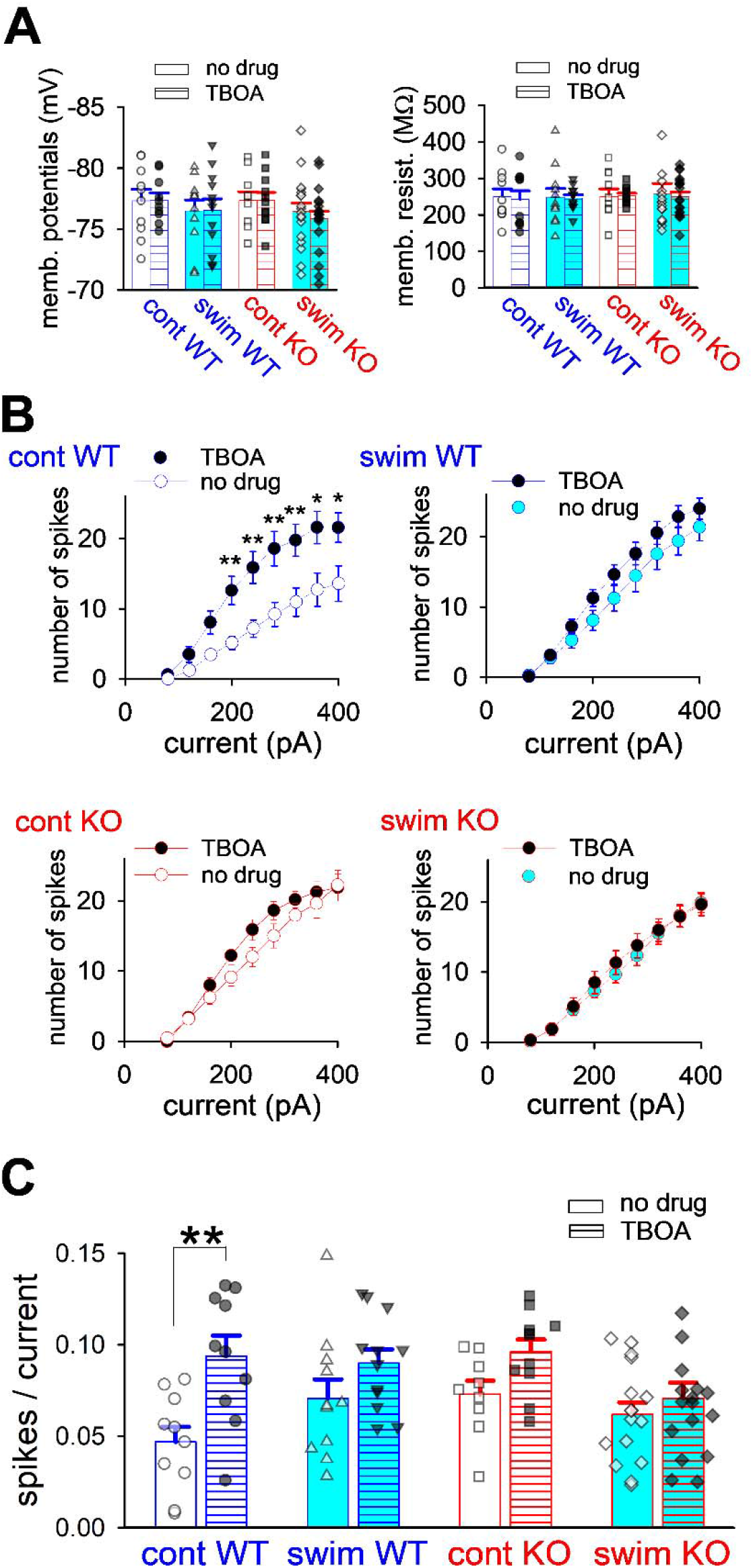
Elevation of spike firing by repeated swim and PKN1a knockout is mimicked and occluded by glutamate transporter inhibition. (A) Summary of the effects of DL-TBOA (100 μM) on resting membrane potentials and membrane resistance (control wild-type, no drug, *n* = 10 from 3 mice, TBOA, *n* = 10 from 3 mice; swim wild-type, no drug, *n* = 11, from 3 mice, TBOA, *n* = 12 from 3 mice; control KO, no drug, *n* = 9 from 3 mice, TBOA, *n* = 11 from 3 mice; swim KO, no drug, *n* = 18 from 3 mice, TBOA, *n* = 19 from 3 mice). (B) Summary of the effects of DL-TBOA on current-spike relationship. *C,* summary of averaged slopes in control and repeated-swim wild-type mice and PKN1a KO mice. \**p* < 0.05. All data are shown as means ± s.e.m.

However, TBOA significantly increased spike firing in control wild-type mice (Fig. 3B; drug, *F*_(1, 18)_ = 8.600, *p* = 0.009; drug × injected current, *F*_(8, 144)_ = 2.838, *p* = 0.006; 200 pA, *p* = 0.007; 240 pA, *p* = 0.002; 280 pA, *p* = 0.001; 320 pA, *p* = 0.002; 360 pA, *p* = 0.016; 400 pA, *p* = 0.035;two-way repeated-measures ANOVA). In contrast, TBOA did not significantly affect spike firing in repeated-swim wild-type (Fig. 3B; drug, *F*_(1, 21)_ = 1.602, *p* = 0.219; drug × injected current, *F*_(8, 168)_ = 1.513, *p* = 0.156; two-way repeated-measures ANOVA), control KO (Fig. 3B; drug, *F*_(1, 18)_ = 1.539, *p* = 0.231; drug × injected current, *F*_(8, 144)_ = 1.255, *p* = 0.272; two-way repeated-measures ANOVA), or repeated-swim KO mice (Fig. 3B; drug, *F*_(1, 31)_ = 0.127, *p* = 0.724; drug × injected current, *F*_(8, 248)_ = 0.443, *p* = 0.894; two-way repeated-measures ANOVA). We also calculated the slope of the relationship between the number of evoked spikes and the injected-current amplitude (Fig. 3C) and found a significant main effect of drug (*F*_(1, 88)_ = 16.161, *p* = 1.22 × 10; three-way ANOVA) and a significant genotype × swim interaction (*F*_(1, 88)_ = 5.323, *p* = 0.0234; three-way ANOVA). TBOA significantly increased slope in in control wild-type mice (Fig. 3C; no drug, 0.0468 ± 0.0083 spikes/pA, TBOA, 0.0938 ± 0.0112 spikes/pA, *p* = 0.0022; Šídák’s multiple-comparisons test), suggesting that depolarization of dentate granule cells increases extracellular glutamate levels and that glutamate transporters normally limit neuronal excitability by clearing extracellular glutamate. In contrast, TBOA did not alter firing in repeated-swim wild-type mice (Fig. 3C; no drug, 0.0706 ± 0.0104 spikes/pA, TBOA, 0.0900 ± 0.0075 spikes/pA) or in either control or repeated-swim PKN1a KO mice (Fig. 3C; control, no drug, 0.0729 ± 0.0074 spikes/pA, control, TBOA, 0.0963 ± 0.0066 spikes/pA; swim, no drug, 0.0618 ± 0.0066 spikes/pA, swim, TBOA, 0.0708 ± 0.0085 spikes/pA). These results suggest that dentate granule cell excitability has already reached a ceiling under these conditions, such that TBOA cannot further enhance spike firing. Thus, repeated swim and PKN1a knockout likely reduce glutamate transporter function, thereby increasing spike firing through enhanced activation of group I mGluRs.

Notably, group I mGluRs appear to be activated to a similar extent at resting membrane potentials in control and repeated-swim wild-type and PKN1a KO mice, because group I mGluR antagonists reduced membrane resistance to a comparable degree under all conditions (Fig. 2A), and glutamate transporters were ineffective at resting membrane potentials in each group (Fig. 3A). Therefore, enhanced tonic activation of group I mGluRs by repeated swim or PKN1a knockout is unlikely, and ambient glutamate concentrations at rest are presumably similar between control and repeated-swim hippocampi in both wild-type and PKN1a KO mice. Taken together, these findings indicate that glutamate transporters efficiently clear glutamate primarily during spike firing in the normal hippocampus, whereas this mechanism is compromised in repeated-swim wild-type and PKN1a KO mice, resulting in elevated neuronal excitability. Glutamate can be released from dendrites and somata and can act as a retrograde or autocrine messenger (Duguid et al., 2007; Harkany et al., 2004; Ovsepian and Dolly, 2011). In addition, glutamate functions as a gliotransmitter (Dallerac et al., 2018; de Ceglia et al., 2023), and depolarization of dentate granule cells may increase extracellular K concentrations, potentially depolarizing astrocytes and promoting astrocytic glutamate release, although the cellular source of glutamate released during neuronal activity remains to be identified.

### 3.2. Repeated swim and PKN1 knockout reduce total and membrane expression of EAAT3

We first examined hippocampal PKN1 expression in wild-type mice subjected to repeated swim because repeated swim increased dentate granule cell excitability and reduced hippocampal EAAT3 expression, phenotypes that were also observed in PKN1a KO mice. However, neither a single swim exposure nor five days of repeated swim significantly altered hippocampal PKN1 expression (Fig. S2; control, n = 12 from 6 mice; 1-day swim, n = 6 from 3 mice; 5-day swim, n = 8 from 4 mice).

Previously, we reported that PKN1a knockout reduces total expression of EAAT3 in the developing hippocampus (Yasuda et al., 2020). Based on this finding, we quantified EAAT3 expression in the hippocampus of PKN1a KO mice and repeated-swim wild-type mice. Analysis of total EAAT3 expression in the hippocampus revealed a significant main effect of repeated swim (Fig. 4A; swim, *F*_(1, 30)_ = 24.613, *p* < 0.001; two-way ANOVA) and a significant genotype × swim interaction (*F*_(1, 30)_ = 23.991, *p* < 0.001; two-way ANOVA), whereas the main effect of genotype was not significant (*F*_(1, 30)_ = 2.615, *p* = 0.116; two-way ANOVA). Total hippocampal EAAT3 expression was significantly reduced in repeated-swim wild-type and control PKN1a KO mice (Fig. 4A; control wild-type, 100 ± 2.8%, *n =* 14 from 7 mice; swim wild-type, 57.9 ± 4.2%, *n =* 8 from 4 mice, *p* < 0.001, post hoc Tukey–Kramer test; control KO, 72.2 ± 4.8%, *n =* 8 from 4 mice, *p* < 0.001; swim KO, 71.9 ± 2.9%, *n =* 4 from 2 mice). In contrast, total hippocampal expression of EAAT2 (also known as glutamate transporter 1, GLT-1), the major glial glutamate transporter that mediates glutamate uptake more efficiently than EAAT3 does (Jensen et al., 2015), was not significantly altered in repeated-swim wild-type mice or in control and repeated-swim PKN1a KO mice (Fig. S4; control wild-type, *n =* 6 from 3 mice, swim wild-type, *n =* 6 from 3 mice; control KO, *n =* 6 from 3 mice, swim KO, *n =* 6 from 3 mice). We next examined surface expression of EAAT3 in the hippocampus using a brain slice biotinylation assay (Nolt et al., 2011). Analysis of surface EAAT3 expression revealed significant main effects of genotype (Fig. 4B; *F*_(1, 24)_ = 19.102, *p* < 0.001) and repeated swim (swim, *F*_(1, 24)_ = 8.820, *p* = 0.007), as well as a significant genotype × stress interaction (*F*_(1, 24)_ = 11.044, *p* = 0.003). Consistent with the changes observed in total protein levels, surface expression of EAAT3 was also significantly reduced in repeated-swim wild-type and in control PKN1a KO mice (Fig. 4B; control wild-type, 100 ± 0.0%, *n =* 9 from 9 mice; swim wild-type, 73.4 ± 5.7%, *n =* 5 from 5 mice, *p* < 0.001; post hoc Tukey–Kramer test, control KO, 70.8 ± 3.7%, *n =* 9 from 9 mice, *p* < 0.001; swim KO, 74.9 ± 5.6%, *n =* 5 from 5 mice). Together, these results indicate that repeated swim and PKN1a knockout selectively reduce both total and surface expression of EAAT3, likely leading to elevated extracellular glutamate levels, enhanced activation of mGluRs, and increased firing of dentate granule cells.

**Figure 4.**
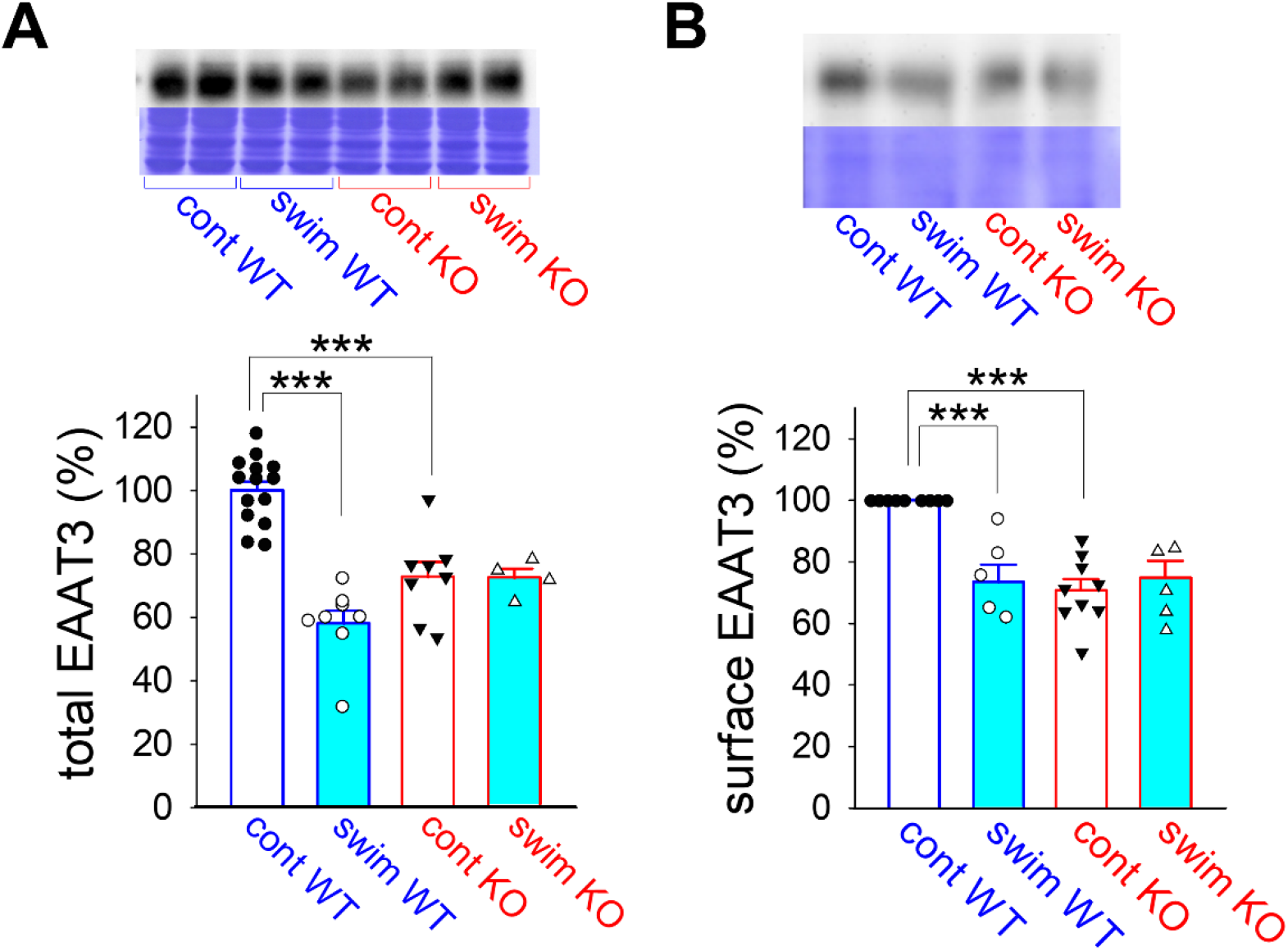
Repeated swim and PKN1a knockout reduce total and surface expression of EAAT3 in mouse hippocampus. (A) Representative EAAT3 immunoblots with corresponding Coomassie-stained protein bands used as loading controls and summary of quantitative immunoblot analyses of total hippocampal EAAT3 expression in control wild-type mice (*n =* 14 from 7 mice), control PKN1a KO mice (*n =* 8 from 4 mice), repeated-swim wild-type mice (*n =* 8 from 4 mice), and repeated-swim PKN1a KO mice (*n =* 4 from 2 mice). (B) Representative EAAT3 immunoblots with corresponding Coomassie-stained protein bands used as loading controls and summary of quantitative immunoblot analyses of surface hippocampal EAAT3 expression in control wild-type mice (*n =* 9 from 9 mice), control PKN1a KO mice (*n =* 9 from 9 mice), repeated-swim wild-type mice (*n =* 5 from 5 mice), and repeated-swim PKN1a KO mice (*n =* 5 from 5 mice). For *B*, the amount of biotinylated EAAT3 in the eluate fraction was quantified. The original uncropped images corresponding to the immunoblot and Coomassie-staining panels are shown in Fig. S3. \*\*\**p* < 0.001. All data are shown as means ± s.e.m.

### 3.3. Repeated swim and PKN1a knockout reduce anxiety-related behavior in elevated plus maze

We have shown that PKN1 normalizes group I mGluR activity by upregulating EAAT3 expression and that repeated swim increases excitability in the dentate gyrus via reduced EAAT3 expression. However, the roles of PKN1 in mouse behavior remain largely unexplored. Repeated swim has been used to induce depression-like behavior in rodents, and increased immobility during swimming has been reported (Boyce-Rustay et al., 2007; Sun et al., 2011). Notably, however, increased immobility in the forced swim test after repeated swim exposure may reflect learned immobility or adaptation to the swim context rather than a depression-like phenotype (Mul et al., 2016). Therefore, we assessed depression-like behavior in repeated-swim and PKN1a KO mice using the tail suspension test. Immobility time did not differ significantly across genotypes or experimental conditions (Fig. S5A; control wild-type, *n =* 12, swim wild-type, *n =* 10; control KO, *n =* 12, swim KO, *n =* 10). Consistent with this, immobility time in the forced swim test did not differ between control wild-type and PKN1a KO mice (Fig. S5B; control wild-type, *n* = 13; control KO, *n* = 12). Together, these results indicate that neither repeated swim nor PKN1a knockout induced depression-like behavior.

Optogenetic activation of the ventral dentate gyrus reduces anxiety (Kheirbek et al., 2013). Based on this finding, we performed the elevated plus maze test to assess anxiety-like behavior. Control PKN1a KO mice spent significantly more time in the open arms than control wild-type mice, indicating reduced anxiety-like behavior in KO mice (Fig. 5A and B; control wild-type, 71.6 ± 9.2 s, *n =* 20; control KO mice, 120.3 ± 10.1 s, *n =* 24, genotype, *F*_(1, 117)_ = 8.95, swim, *F*_(2, 117)_ = 5.85, *p* = 0.004, two-way ANOVA with Tukey’s test). Repeated swim significantly increased the time spent in the open arms in wild-type mice (Fig. 5A and B; 5-day swim, 125.1 ± 12.5 s, *n =* 19, *p* = 0.008; 10-day swim, 120.3 ± 10.9 s, *n =* 26, *p* = 0.018; two-way ANOVA with Tukey’s test), but did not affect open-arm time in PKN1a KO mice (Fig. 5A and B; 5-day swim, 142.6 ± 18.8 s, *n =* 15; 10-day swim, 138.7 ± 13.6 s, *n =* 19). Together, these results suggest that PKN1a expression promotes cautious behavior under baseline conditions, whereas exposure to mild stress, such as repeated swim, shifts behavior toward reduced anxiety-like responses in potentially threatening environments.

**Figure 5.**
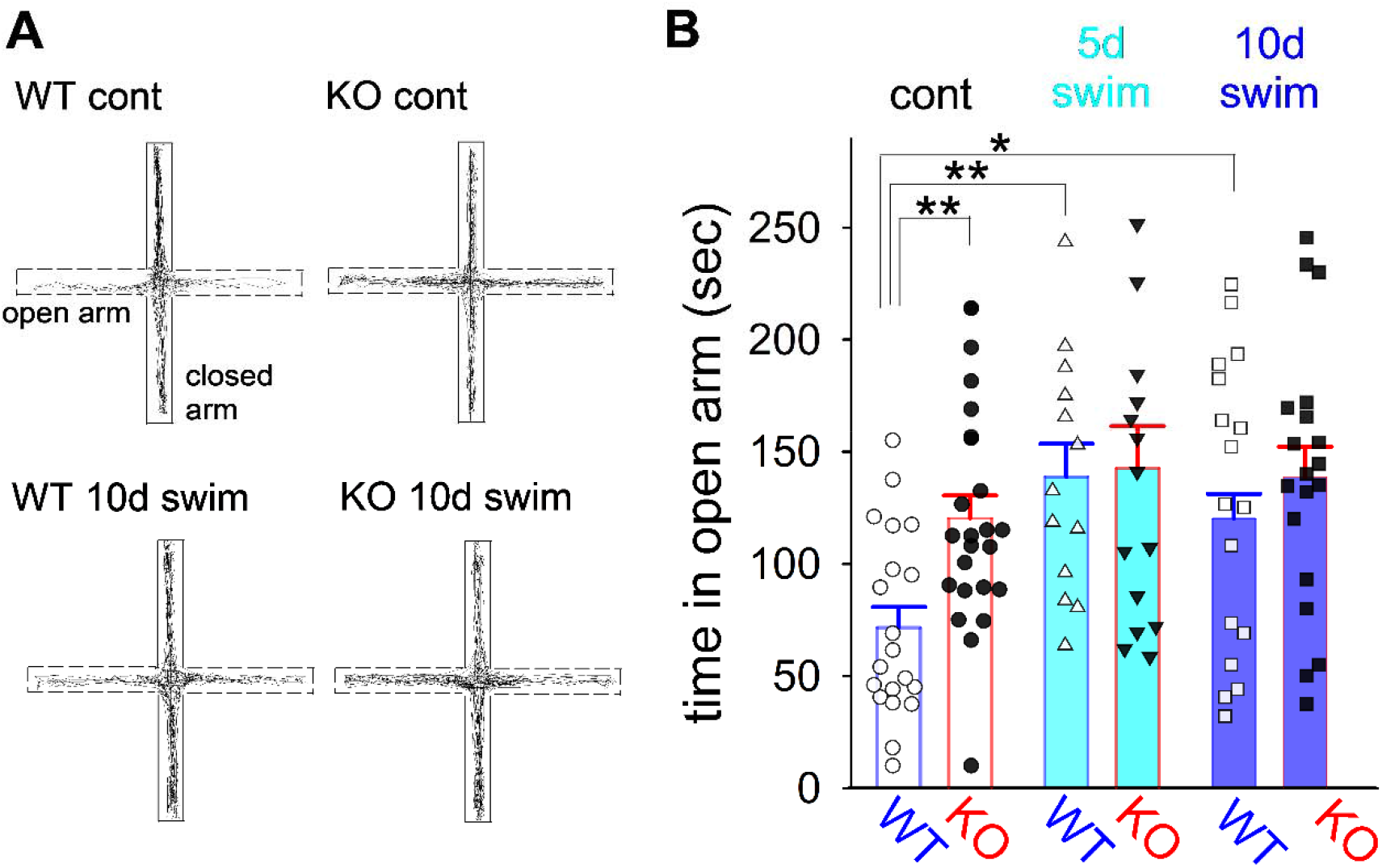
PKN1a knockout and repeated swim reduce anxiety-related behavior in elevated plus maze test. (A and B) Track traces (A) and summary (B) of time spent in the open arms in the elevated plus maze test. Time spent in the open arms was longer in PKN1a KO mice (*n =* 24) than in wild-type mice (*n =* 20) under control conditions. Although repeated swim did not alter time spent in the open arms in PKN1a KO mice (5-day swim, *n =* 15; 10-day swim, *n =* 19), repeated swim increased time spent in the open arms in wild-type mice (5-day swim, *n =* 19; 10-day swim, *n =* 26). \**p* < 0.05, \*\**p* < 0.01. All data are shown as means ± s.e.m.

## 4. Discussion

Group I mGluRs mediate LTD and induce neuronal depolarization and have been implicated in various brain disorders, including anxiety, depression, and addiction (Luscher and Huber, 2010; Niswender and Conn, 2010; Pilc et al., 2008). However, the mechanisms that regulate their activity remain poorly understood. In the present study, we found that both PKN1a knockout and repeated swim reduced total and surface EAAT3 expression and were associated with enhanced group I mGluR-dependent activity and increased spike firing in mature dentate granule cells of the mouse hippocampus. Furthermore, PKN1a knockout and repeated swim reduced anxiety-related behaviors in the elevated plus maze test. These findings suggest that PKN1a contributes to the maintenance of cautious behavior under baseline conditions; however, following mild stress exposure, wild-type mice show reduced anxiety-related behavior in elevated environments.

### 4.1. PKN1a-dependent regulation of EAAT3 expression helps restrain group I mGluR-dependent excitability

PKN1 increases both total and surface expression of the neuronal glutamate transporter EAAT3, thereby reducing group I mGluR activity by lowering extracellular glutamate concentrations and suppressing firing in dentate granule cells. EAAT3 is modestly expressed in hippocampal neurons and has a lower capacity for glutamate clearance than the glial transporter EAAT2 (Jensen et al., 2015). Although glial EAAT2 expression is reduced in rats with learned helplessness and glutamate clearance from the extracellular space is decreased in multiple brain regions, including the hippocampus (Popoli et al., 2012), repeated swim and PKN1a knockout did not significantly alter EAAT2 expression in the mouse hippocampus (Fig. S4). However, even a modest reduction of surface EAAT3 by PKN1a knockout and repeated swim (Fig. 4B) enhanced group I mGluR-dependent firing in mature dentate granule cells (Fig. 2). A similarly small reduction in EAAT3 expression also induces group I mGluR–dependent LTD in the CA1 region of the developing mouse hippocampus (Yasuda et al., 2020). These results suggest that glutamate uptake through neuronal EAAT3 is relatively more important, and that glial glutamate uptake contributes less to regulation of extracellular glutamate concentration in the hippocampus than in other brain regions. Many synapses are not covered by glial processes in the hippocampus (Ventura and Harris, 1999); therefore, EAAT3 is likely to take up glutamate at hippocampal synapses more effectively than glial EAAT2 and downregulate synaptic mGluR activity (Jarzylo and Man, 2012; Yasuda et al., 2020). Fewer astrocytic cell bodies and processes are present in the cell layers than in other layers of the mouse CA1 region and dentate gyrus (Kim et al., 2018; Viana et al., 2023). Therefore, dentate granule cell bodies may not be efficiently covered by astrocytic processes, and glutamate uptake by astrocytes may be insufficient, such that even a modest decrease in neuronal EAAT3 expression affects firing in dentate granule cells. Repeated swim did not alter hippocampal PKN1 expression (Fig. S2), arguing against a simple mechanism in which repeated swim reduces EAAT3 expression by decreasing PKN1 abundance. Nevertheless, an effect of repeated swim on PKN1 activity cannot be excluded. PKN1 is activated by arachidonic acid (Mukai, 2003), whereas steroid hormones and stress mediators can inhibit phospholipase A2, which releases arachidonic acid from phospholipids (Farooqui et al., 2007). Thus, stress could potentially modulate PKN1 activity by altering arachidonic acid availability. However, PKN1 activity was not directly examined in the present study, and our data do not establish PKN1a as a causal mediator of the reduction in EAAT3 expression induced by repeated swim. At present, PKN1a knockout and repeated swim should therefore be considered distinct genetic and environmental manipulations that converge on reduced EAAT3 expression and enhanced group I mGluR-dependent excitability. Accordingly, PKN1a knockout provides a genetic means of identifying a PKN1a-dependent mechanism regulating EAAT3 expression and granule cell excitability, rather than serving as a genetic model of repeated swim exposure.

Repeated stress alters neuronal excitability in a brain region-dependent manner, and the involvement of K^+^ channels has been suggested (Russo et al., 2012). Chronic social isolation upregulates several K^+^ channels and lowers neuronal excitability in the nucleus accumbens (Wallace et al., 2009). Forced swim stress increases spike firing with spike broadening by inactivating large-conductance Ca^2+^-dependent BK channels in the cingulate cortex (Sun et al., 2011). In contrast, BK channel dysfunction results in spike broadening and reduced spike firing (Brenner et al., 2005). We found that repeated swim and PKN1a knockout increased spike firing without changes in spike width in mature dentate granule cells (Fig. S1A-D), suggesting that repeated swim and PKN1a knockout have minimal effects on calcium-dependent K^+^ channel activity. In addition, voltage responses to injected currents in the presence of TTX and Cd^2+^ did not differ between control and repeated-swim wild-type mice or between control and repeated-swim PKN1a KO mice (Fig. S1E). Therefore, changes in K^+^ channels are unlikely to account for elevated spike firing in repeated-swim and PKN1a KO mice. Rather, our results suggest that neuronal glutamate transporter function is reduced by repeated swim or PKN1a knockout, and that enhanced group I mGluR activation leads to increased spike firing in dentate granule cells, because the elevated spike firing induced by repeated swim or PKN1a knockout was suppressed by group I mGluR antagonists (Fig. 2) and was mimicked and occluded by a glutamate transporter inhibitor DL-TBOA (Fig. 3).

Although the downstream ionic mechanisms linking group I mGluR activation to increased spike firing were not directly examined in the present study, several candidate mechanisms have been reported in the hippocampus. Group I mGluR activation suppresses a Ca²-activated K current in dentate granule cells (Abdul-Ghani et al., 1996) and enhances neuronal excitability by activating voltage-dependent inward currents and TRPC-mediated nonselective cation currents and by enhancing Nav1.6-mediated persistent Na currents in the hippocampus (Chuang et al., 2000; El-Hassar et al., 2011; Yu et al., 2018). These mechanisms could preferentially enhance excitability during depolarization without necessarily altering the resting membrane potential. Whether any of these mechanisms mediate the group I mGluR-dependent increase in dentate granule cell excitability observed in the present study remains to be determined.

### 4.2. Repeated swim exposure and PKN1a knockout are associated with reduced anxiety-related behavior

Repeated swim has been used to induce depression-like behavior, and increased immobility during swimming has been reported (Boyce-Rustay et al., 2007; Sun et al., 2011), although this immobility could simply reflect habituation of mice to water, and its ability to induce depression-like behavior is debatable (Mul et al., 2016). The effects of repeated swim on anxiety are also controversial. Repeated swim induces anxiety-like behavior in Sprague-Dawley rats (Avital et al., 2001); however, it does not induce anxiety-like behavior in mice (Boyce-Rustay et al., 2007; Mul et al., 2016). Unexpectedly, 5 and 10 days of repeated swim reduced anxiety-related behavior in wild-type mice in the elevated plus maze test (Fig. 5), suggesting that repeated swim may act less as a stressor and more as a mild exercise-like challenge that enhances stress resilience. Repeated swim is also used as an exercise in rodents (Sun et al., 2021; Yu et al., 2024). Repeated swim for 15–60 min per day for 5 weeks does not induce anxiety– or depression-like behavior; however, it prevents anxiety– and depression-like behavior induced by chronic unpredictable mild stress (Sun et al., 2021). A similar repeated swim protocol (20–60 min daily for 4 weeks) also improves cardiac function, reduces blood pressure, and blood glucose levels, even though it increases body weight (Yu et al., 2024). Exercise has both anxiolytic (Shen et al., 2024; Yan et al., 2024) and anxiogenic effects (Fuss et al., 2010). We presume that the anxiolytic effects of repeated swim may be mediated by activation of most of the muscles in the mouse body during swimming. Such extensive muscle contractions may engage exercise-associated peripheral mechanisms, including the release of myokines that can influence brain function and promote stress resilience (Liu et al., 2026; Yu et al., 2026). Thus, unlike restraint or social defeat, repeated swim may have both an aversive stress component and an exercise-related adaptive component. We therefore speculate that, under the conditions used in the present study, the balance between these components may have favored the latter, potentially leading to the recruitment of processes or neural circuits that suppress anxiety-related behavior and thereby contributing to the increased open-arm time in the elevated plus maze.

An important limitation of the present study is that the use of non-swum animals as controls does not allow us to determine the relative contributions of the aversive and exercise-related components of repeated swim exposure. An ideal control would reproduce the amount and pattern of swimming-associated whole-body muscular activity without water immersion, inescapability, or other aversive aspects of the procedure. However, establishing such a control is difficult. Voluntary wheel running or treadmill exercise would differ from swimming in terms of controllability, sensory environment, and the pattern of muscle recruitment, whereas shallow-water exposure would not reproduce the muscular activity associated with swimming. Thus, no single comparison group would be expected to match all aspects of repeated swim exposure other than its aversive component. Previous repeated-swim studies have commonly used non-swum or home-cage animals as controls (Baganz et al., 2010; Boyce-Rustay et al., 2007; Kiselycznyk et al., 2011; Mul et al., 2016), as in the present study. Accordingly, our results demonstrate the overall effects of repeated swim exposure but do not distinguish whether these effects are mediated by its aversive component, exercise-related component, or an interaction between the two.

We found that repeated swim exposure and PKN1a knockout increased spike firing through enhanced group I mGluR activity in dentate granule cells (Fig. 2) and reduced anxiety-related behavior in the elevated plus maze test (Fig. 5). These findings raise the possibility that selective enhancement of group I mGluR signaling in dentate granule cells may increase their activity and reduce anxiety-related behavior. Nevertheless, although pharmacological modulation of group I mGluRs has yielded promising preclinical results, clinical efficacy has not been consistently demonstrated for either depression (Dogra and Conn, 2021; Quiroz et al., 2016) or anxiety disorders (Dogra and Conn, 2021; Ferraguti, 2018). One possible explanation for this translational gap is that the behavioral consequences of group I mGluR modulation depend on the brain region and neural circuit involved. For example, systemic administration of group I mGluR antagonists produces anxiolytic-like effects (Niswender and Conn, 2010; Pilc et al., 2008), whereas reduced mGluR5 signaling in the dorsal hippocampal CA1 region induces anxiety-related behavior that is alleviated by local administration of an mGluR5 positive allosteric modulator (Li et al., 2023). Together with our findings, these observations suggest that either enhancement or inhibition of group I mGluR signaling may reduce anxiety-related behavior depending on the neural circuit involved. Moreover, clinical development of group I mGluR modulators has largely focused on mGluR5, although mGluR1 antagonists have also been reported to produce anxiolytic-like effects in rodents (Dogra and Conn, 2021). In our previous study, enhanced mGluR-dependent LTD in PKN1a KO mice involved both mGluR1 and mGluR5 (Yasuda et al., 2020). Therefore, in pathological circuits in which both receptor subtypes contribute to excessive group I mGluR signaling, inhibition of mGluR5 alone may be insufficient, and simultaneous inhibition of mGluR1 and mGluR5 might produce greater anxiolytic efficacy. This possibility warrants further investigation.

As mentioned previously, overexpression of EAAT3 in the mouse forebrain increases anxiety-related behavior (Delgado-Acevedo et al., 2019), and maternal separation reduces hippocampal EAAT3 expression and anxiety-related behavior (Kim et al., 2020). In line with these results, PKN1a knockout and repeated swim reduced total and surface expression of EAAT3 in the mouse hippocampus and reduced anxiety-related behavior. Reduced EAAT3 expression is associated with elevated spike firing in the dentate gyrus. Optogenetic activation of the ventral dentate gyrus reduces anxiety-related behavior (Kheirbek et al., 2013). Therefore, elevated excitability in dentate granule cells in PKN1a KO and repeated-swim wild-type mice may contribute to the reduction of anxiety-related behavior. In addition, recent studies suggest that elevated EAAT3 expression makes mice less susceptible to chronic stress that can induce depression-like behavior (Ardiles et al., 2025), whereas reduced EAAT3 expression in the rodent hippocampus is associated with depression-like behavior (Kim et al., 2020). However, PKN1a knockout and repeated swim did not induce depression-like behavior (Fig. S4), presumably because the decreases in EAAT3 expression in PKN1a KO and repeated-swim mice were not as large as those in EAAT3 KO mice or maternally separated mice (Kim et al., 2020). EAAT3 has also been reported to be involved in several human brain disorders, including bipolar disorder, epilepsy, obsessive–compulsive disorder, and schizophrenia (Bianchi et al., 2014; Bjorn-Yoshimoto and Underhill, 2016; Delgado-Acevedo et al., 2019). In contrast, the involvement of PKN1 in brain disorders has been less extensively characterized. PKN1 accumulates in neurofibrillary tangles and phosphorylates tau in Alzheimer’s disease (Kawamata et al., 1998). In addition, PKN1-N207, a truncated peptide generated by inclusion of the TDP-43–repressed cryptic exon PKN1-5a1, has been detected in Alzheimer’s disease brains with TDP-43 pathology (Yang et al., 2026). Despite these findings, the broader roles of PKN1 in neurological and neuropsychiatric disorders remain largely unclear. The present study identified a link between PKN1a-dependent regulation of EAAT3 expression and group I mGluR-dependent excitability in dentate granule cells, but did not establish whether this mechanism contributes causally to anxiety-related behavior. Future in vivo studies using local pharmacological manipulation of group I mGluRs or EAAT-mediated glutamate uptake in the hippocampal dentate gyrus will help clarify whether the PKN1a–EAAT3–group I mGluR pathway contributes to the regulation of anxiety-related behavior.

## Data Availability

The data generated in this study are available from the corresponding author upon reasonable request.

## CRediT authorship contribution statement

**Hiroki Yasuda:** Conceptualization, Project administration, Supervision, Funding acquisition, Methodology, Resources, Investigation, Data curation, Formal analysis, Validation, Visualization, Writing – original draft, Writing – review & editing. **Koji Kubouchi:** Writing – review & editing, Investigation. **Kenji Hanamura:** Writing – review & editing, Investigation, Methodology. **Taiga Kurihara:** Writing – review & editing, Investigation. **Yuka Nakasone:** Writing – review & editing, Investigation. **Hideyuki Mukai:** Writing – review & editing, Visualization, Investigation, Methodology, Funding acquisition, Conceptualization, Investigation, Resources.

## Declaration of Generative AI and AI-assisted technologies in the writing process

During the preparation of this work, the authors used Paperpal and ChatGPT to assist with text drafting and editing. After using these tools, the authors reviewed and edited the content as needed and take full responsibility for the content of the manuscript.

## Acknowledgements

We thank Dr. M. Dalva for providing the slice biotinylation protocol.

