## Supplementary Information for "Repeated swim exposure and PKN1a knockout enhance group I mGluR-dependent excitability associated with reduced EAAT3 expression in mouse dentate granule cells"

**Supplementary material**

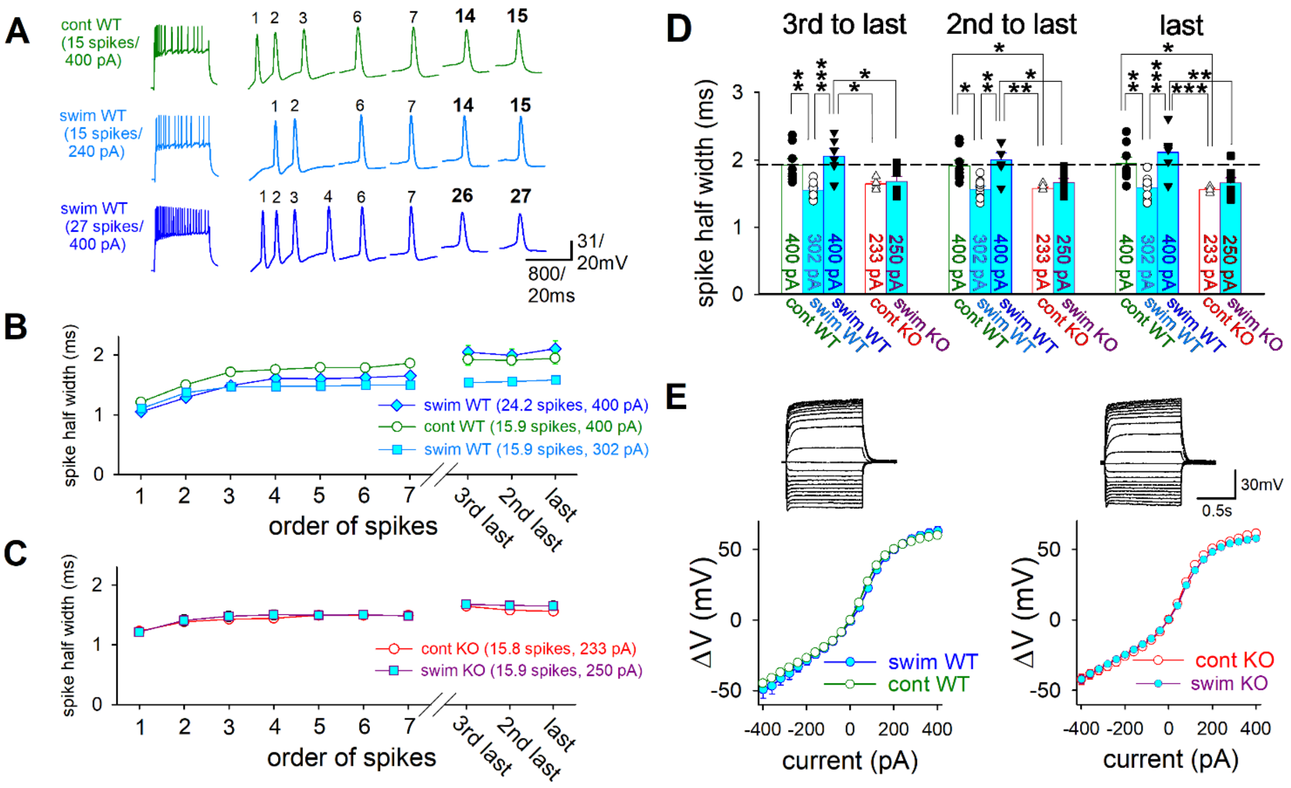

**Figure S1. Minimal involvement of $Ca^{2+}$- or voltage-dependent $K^+$ channels in elevated firing in repeated-swim or PKN1a-deleted dentate granule cells.**

(A) Examples of a 15-spike train induced by a current injection with a maximum current amplitude of 400 pA (1 s) in a control wild-type cell (upper, green), a train with the same number of spikes induced by a smaller current injection (240 pA) in a repeated-swim wild-type cell (middle, cyan), and a 27-spike train induced by the same 400 pA current injection in the repeated-swim wild-type cell (bottom, blue). (B) Summary of spike half-width in spike trains induced by 400 pA current injection in control and repeated-swim wild-type mice (control, 15.9 ± 1.3 spikes, $n$ = 8 from 2 mice (green); repeated swim, 24.2 ± 1.6 spikes, $n$ = 6 from 2 mice (blue)). Trains with 14–16 spikes in repeated-swim wild-type mice were chosen so that the average number of evoked spikes was similar to that in control wild-type mice, and the spike half-width is also shown (15.9 ± 0.7 spikes, average current injection amplitude, 302.2 ± 19.0 pA, $n$ = 9 from 2 mice (cyan)). (C) Summary of spike half-width in control and repeated-swim PKN1a KO mice (control, 15.8 ± 0.3 spikes, average current injection amplitude, 233.3 ± 21.7 pA, $n = 6$ from 3 mice (red); repeated swim, 15.9 ± 0.4 spikes, average current injection amplitude, 250.0 ± 12.5 pA, $n = 8$ from 3 mice (dark pink)). Repeated swim did not broaden spikes in wild-type and KO mice. (D) Summary of half-widths of the third-from-last, second-from-last, and last spikes in control and repeated-swim wild-type and PKN1a KO mice. (E) Voltage responses to injected currents in the presence of $Na^+$ and $Ca^{2+}$ channel inhibitors (TTX 1 μM, $Cd^{2+}$ 200 μM) did not differ between control and repeated-swim wild-type (control, $n = 11$ from 2 mice; swim, $n = 9$ from 2 mice) and PKN1a KO mice (control, n = 11 from 2 mice; swim, n = 16 from 2 mice), suggesting that $K^+$ conductance was not significantly altered by repeated swim or PKN1a knockout. *$p < 0.05$, **: $p < 0.01$, ***$p < 0.001$. All data are shown as means ± s.e.m.

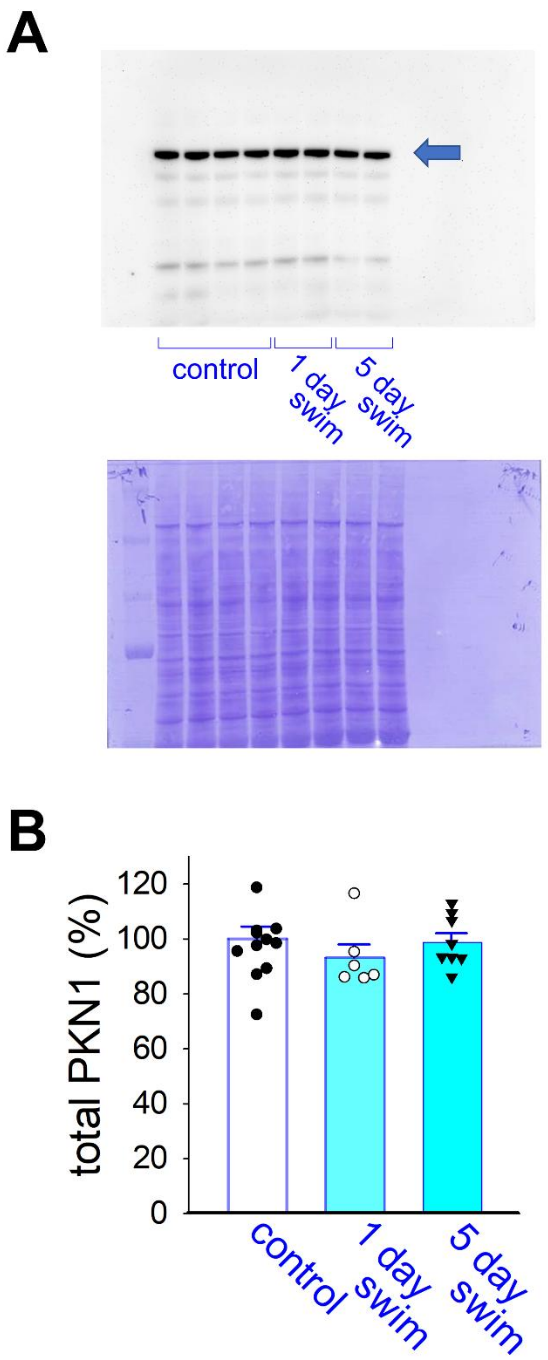


**Figure S2. Repeated swim does not change total expression of PKN1 in wild-type mouse hippocampus.** (A and B) Representative PKN1 immunoblots (arrow) with corresponding Coomassie-stained protein bands used as loading controls (A), and summary (B) of quantitative immunoblot analyses of hippocampal PKN1 expression in control (n = 12 from 6 mice), one-day swim (*n* = 6 from 3 mice), and five-day swim wild-type mice (n = 8 from 4 mice).

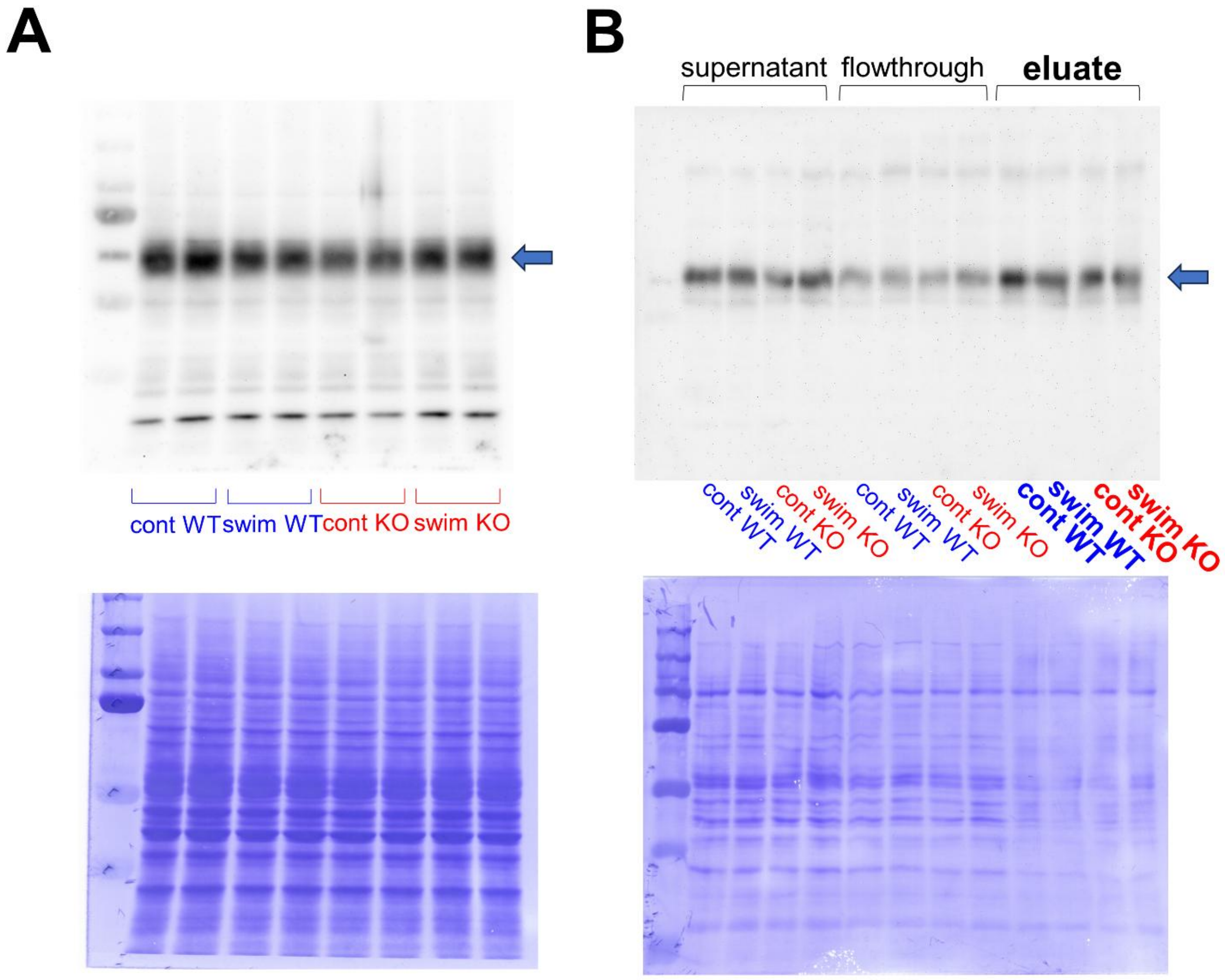


**Figure S3. Uncropped immunoblot images shown in Fig. 4.**

(A) Original immunoblot images shown in Fig. 4A and Coomassie staining of the same membranes. (B) Original immunoblot images shown in Fig. 4B and Coomassie staining of the same membranes.

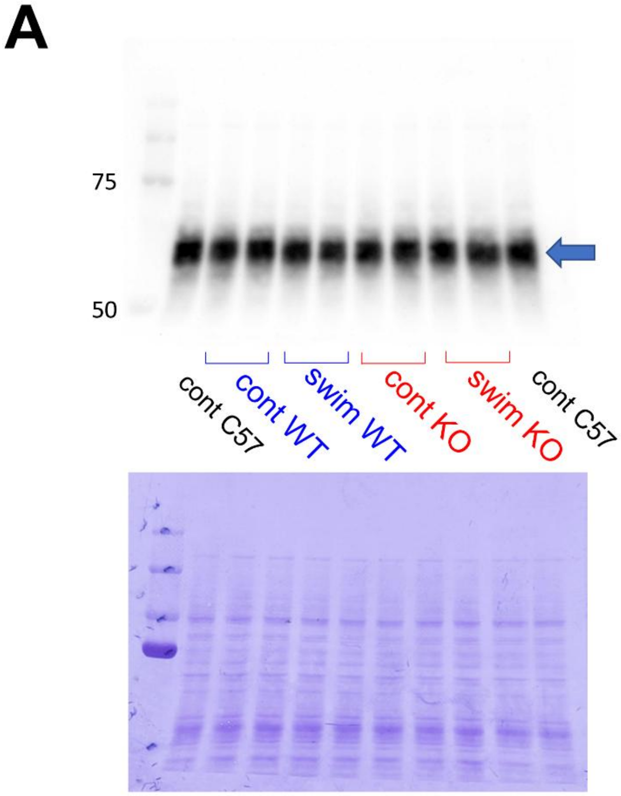


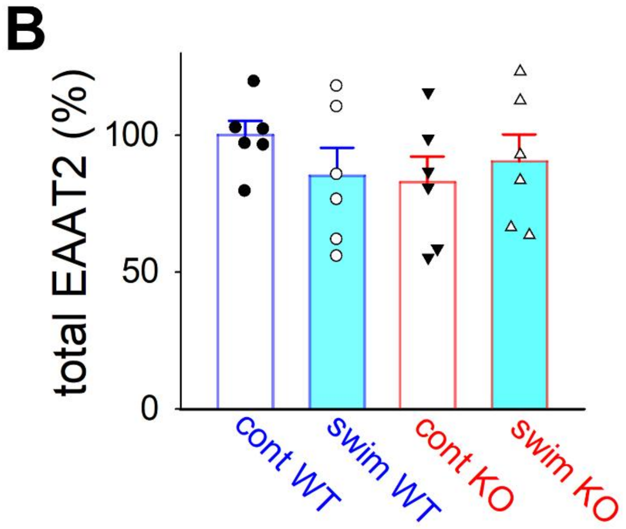


**Figure S4. Repeated swim and PKN1a knockout do not change total expression of EAAT2 in mouse hippocampus.**

(A and B) Representative EAAT2 immunoblots (arrow) with corresponding Coomassie-stained protein bands used as loading controls (A), and summary (B) of quantitative immunoblot analyses of hippocampal EAAT2 expression in control wild-type mice (*n* = 6 from 3 mice), repeated-swim wild-type mice (*n* = 6 from 3 mice), control PKN1a KO mice (*n* = 6 from 3 mice), and repeated-swim PKN1a KO mice (*n* = 6 from 3 mice).

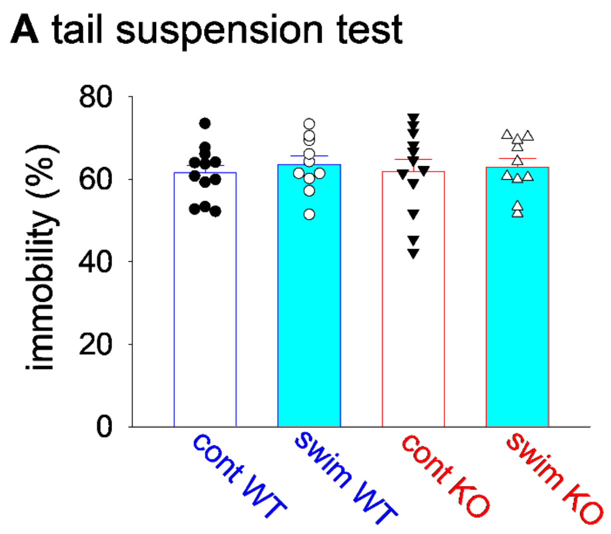


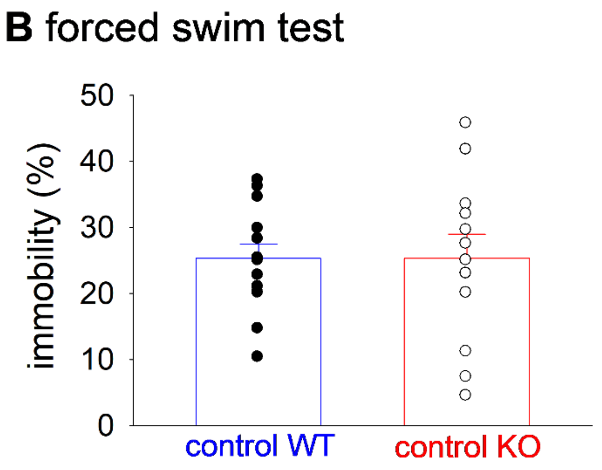


**Figure S5. Repeated swim and PKN1a knockout do not alter immobility time in tail suspension and forced swim tests.**

(A) Summary of immobility time in control wild-type mice (n = 12), repeated-swim wild-type mice (n = 10), control PKN1a KO mice (n = 12), and repeated-swim PKN1a KO mice (n = 10) in the tail suspension test. (B) Summary of immobility time in control wild-type mice (n = 13) and PKN1a KO mice (n = 12) in the forced swim test.
